# Structural and functional characterisation of the Crimean-Congo Haemorrhagic Fever Virus RNA Dependent RNA Polymerase

**DOI:** 10.1101/2025.10.21.683639

**Authors:** Adrian Deng, Rory Cunnison, Loïc Carrique, Franziska Günl, Els Paddon, Jan Staeyart, David Nguyen Duc, Jonathan M. Grimes, Nicole Robb, Jeremy R. Keown

**Affiliations:** School of Life Sciences, University of Warwick, Coventry, United Kingdom; Warwick Medical School, University of Warwick, Coventry, United Kingdom; Division of Structural Biology, Centre for Human Genetics, University of Oxford, Oxford, United Kingdom; Sir William Dunn School of Pathology, University of Oxford, Oxford, United Kingdom; Structural Biology Brussels, Vrije Universiteit Brussel, VUB, Brussels, Belgium; VIB-VUB Center for Structural Biology, VIB, Brussels, Belgium

**Keywords:** . RNA virus, viral polymerase, bunyavirus, CCHFV, L-protein

## Abstract

Crimean-Congo Haemorrhagic Fever Virus (CCHFV) is found across Africa, Asia, and the Middle East where it can cause Haemorrhagic outbreaks with high case fatality rates. Central to the viral life cycle is the viral L-protein, a crucial and multifunctional protein which both transcribes and replicates the viral genome. Here, we present the cryoEM structures of an RNA free and a 5’ promoter bound complex, describing the core catalytic RNA-dependent RNA polymerase (RdRp). We observe an RdRp that is substantially larger than related L-proteins and contains domain insertions unique to the nairovirus family. The 5’ RNA promoter is found in a tight RNA hairpin stabilised by a single base pair, with 5’ binding triggering the closure of protein over the RNA. Functional analysis of the endonuclease and RdRp activities reveals an enzyme which is capable of both activities and demonstrate RdRp inhibition by known antiviral nucleosides. These data advance our understanding of the molecular mechanisms behind genome replication and transcription, that will help inform future antiviral development.

## Introduction

Crimean-Congo Haemorrhagic (CCHFV, *Orthonairovirus haemorrhagiae*) belongs to the family *Nairoviridae* within the *Bunyaviricetes* class which contains several emerging human and animal pathogens^1^. The Nairovirus family contains over sixty species, with CCHFV causing the majority of human infections and a smaller number of infections being caused by Nairobi sheep disease virus (NSDV)^2^. CCHFV is a tick-borne arbovirus with a broad geographic distribution that includes Africa, the Middle East, Asia and, more recently, southern and eastern Europe^3^. Human infection can lead to haemorrhagic disease with case fatality rates exceeding 30%^4^, although the true incidence is likely underestimated due to a high proportion of subclinical infections. The virus is primarily transmitted by ticks from the *Hyalomma* genus^5^, but human infections can occur through contact with infected animal material.

CCHFV genome organisation is similar to other *Bunyaviricetes,* comprising a negative sense segmented RNA genome^1^. The genome segments are called small (S), medium (M), and large (L). The S segment encodes the viral nucleoprotein (NP) and in an opposite sense open reading frame the small non-structural (NSs) protein^6^. The M segment is more complex than typical *Bunyaviricetes,* encoding not only the viral glycoproteins (Gn and Gc) which perform receptor entry and binding, but also the GP160/85 precursor protein, and a medium non-structural protein (NSm). The L segment encodes the CCHFV L-protein (CCHFV-L). In addition to the coding region, each segment, possesses short untranslated regions at the 5’ and 3’ termini^7^.

Each negative sense RNA segment is packaged into viral ribonucleoprotein (vRNP) complexes that contain a single RNA segment, many copies of NP which bind and protect the RNA, and a single copy of CCHFV-L. The vRNP are the structural and functional units of genome transcription and replication with CCHFV-L providing the enzymatic activities to perform these processes. Transcription initiates with ‘cap-snatching’, a process where CCHFV-L binds host mRNA caps, cleaves a short capped RNA primer, which is then annealed to the 3’ termini of the genome and extended. Genome replication proceeds in a two-step process. First a positive sense complimentary RNA (cRNA) intermediate is synthesised, which is used as a template to produce new vRNA copies in the second step^8^.

Recently L-protein structures have been elucidated for several members of the *Bunyaviricetes* class. These include members of the *Phenuiviridae*^9–12^, *Arenaviridae*^13,14^, *Peribunyaviridae*^15–17^, *Tospoviridae*^18^, and *Hantaviridae*^19–22^. The L-proteins from these viral families are single polypeptides of approximately 250 kDa that contain endonuclease, RNA-dependent RNA polymerase (RdRp), and cap-binding domains^8^. These structures present snapshots of L-proteins without RNA and in conformations related to discrete stages of replication and transcription, mapping the location of the 5’ promoter binding site and the 3’ RNA in several positions.

In contrast to the properties of the viral L-proteins described above, CCHFV-L is almost twice the size at approximately 450 kDa. The N-terminal 169 amino acids of the polypeptide encode an Ovarian tumour like (OTU) protease domain for which several structures have been determined alone and in complex with the interferon-stimulated gene 15 (ISG15) or ubiquitin^23–25^. Residues 587-895 in CCHFV-L encode a putative endonuclease domain with D693 as a key active site residue as identified by virus like particle assays^26^. In vitro analyses of the purified endonuclease domain showed thermal stabilisation in the presence of divalent metal ions or endonuclease inhibitors, but no cleavage activity^27^. The structure of the RdRp has been predicted using pre-AlphaFold *in silico* methods, though the confidence of these models and their utility in antiviral development remained unclear^28^. Functional studies on the activity of the RdRp domain have identified D2517 as the key active site residue, likely coordinating divalent metal ions for NTP hydrolysis^29^. Polymerase activity was demonstrated using a single promoter RNA mimicking the 3’ end of the RNA template and a short complimentary primer and efficient inhibition of the activity by ribavirin or favipiravir^29^. While CCHFV-L is expected to contain a cap-binding domain in the C-terminal region this has not been identified.

In this study we sought to provide a characterisation of the molecular biology of the full-length CCHFV-L. We established robust expression and purification, allowing us to produce both wildtype and several mutant constructs. Initial assessment of the endonuclease activity of the full-length protein revealed a highly active domain, with low metal ion specificity. Guided by structure prediction and previous reports, residues were identified for mutagenesis to generate less active variants for downstream experiments. Single particle electron cryo-microscopy (cryoEM) was subsequently used to determine RNA free and 5’ promoter bound structure at high resolution. These models describe an RdRp domain of approximately 2200 amino acids in length and a unique arrangement of the 5’ promoter binding site. Functional characterisation reveals CCHFV-L to require the 5’ promoter bound for efficient production of RNA products. Collectively, our findings provide a detailed characterisation of a key enzyme from a potent human pathogen to support future antiviral development.

## Results

### Purification and solution characterisation of the CCHFV-L

The gene encoding the full length *Orthonairovirus haemorrhagiae (*Crimean-Congo Haemorrhagic Fever Virus, CCHFV) CCHFV-L with an N-terminal OctaHis tag, a twinStrep, and a TEV-protease site was synthesised and cloned into the pFastbac vector for expression in SF9 insect cells. The protein was purified by affinity capture with SDS-PAGE analysis showing a pure protein sample (Fig 1a). Analysis of the CCHFV-L solution state showed most particles had a mass matching that of monomeric CCHFV-L (Fig 1b) (452.9 kDa from sequence). Additional species at 252 kDa and 840 kDa were also observed, potentially representing small populations of degraded proteins and oligomeric species, respectively.

**Figure 1.**
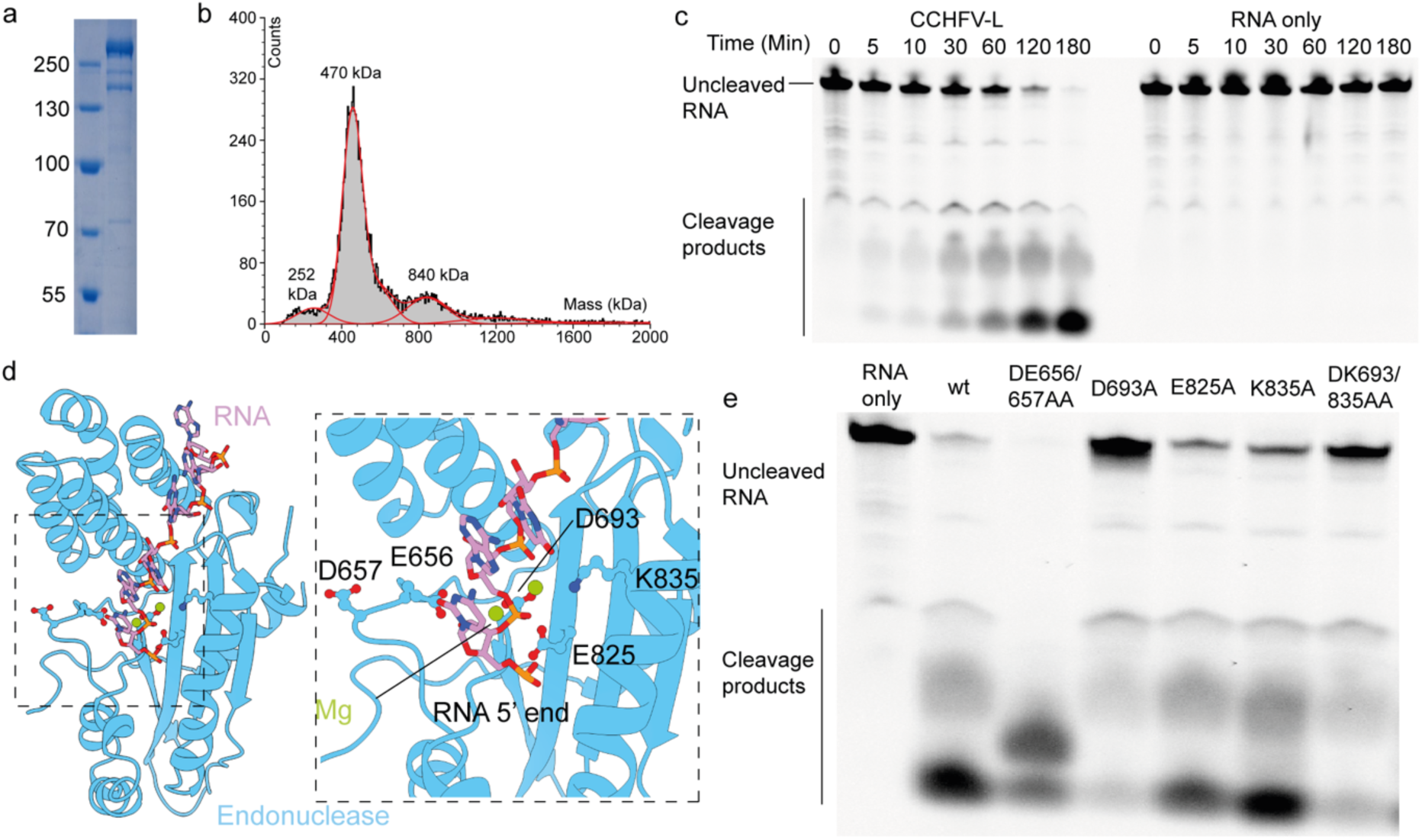
Full-length CCHFV-L contains an active endonuclease domain. a) SDS-PAGE analysis of purified CCHFV-L. b) Mass photometry analysis of full length CCHFV-L. c) Time course RNA degradation of a fluorescently labelled RNA template in the presence or absence of wild type (wt) CCHFV-L. d) Alphafold 3 prediction of a CCHFV endonuclease domain (blue) in complex with magnesium (green) and a segment of RNA (pink). e) Panel of the RNA degradation assay showing RNA products after incubation with CCHFV-L endonuclease proteins for 120 minutes.

### CCHFV-L Endonuclease activity

As previous studies have reported conflicting observations regarding the endonuclease activity of CCHFV-L, we first assessed the activity of our CCHFV-L preparation. A 25 nt fluorescently labelled RNA substrate was incubated with CCHFV-L, and samples were collected at time points between 0-180 minutes (Fig 1c). This experiment showed efficient endonuclease activity, highlighting RNA degradation to short fragments within 120 minutes. No RNA degradation was observed when CCHFV-L was omitted.

Sequence alignments of nairovirus endonuclease domains have suggested residues that are potentially important for activity^27^. To validate and expand the candidate list of important residues in the active site, we performed in silico modelling of the CCHFV endonuclease domain (residues 587-895) in complex with two magnesium ions and a 17 nt RNA fragment. The generated models showed high confidence in the overall domain architecture, confidently predicting the core structure but showing low confidence for two long insertion loops (Supplementary figure 1a). The residues that coordinate the active site magnesium ions and the ions themselves were predicted with high confidence (Supplementary figure 1b). The central portion of the RNA was predicted with moderate confidence, though this was sufficient to identify candidate residues for mutagenesis aimed at reducing endonuclease activity (Fig 1d). Residues E656, D657, D693, E825, and K835 were mutated to alanine and then tested for endonuclease activity. Each mutant protein was assessed for cleavage after 120 minutes of incubation with the fluorescent RNA (Fig 1e). Endonuclease activity was markedly decreased for mutants D693A and K835A, while others showed only little changes. In the predicted model both D693 and K835 directly coordinate the active site magnesium ion. Notably, the D693A mutant has been previously reported to inhibit mRNA production using a minigenome reporter system^26^. The DD656/657AA double mutant appeared to enhance activity. Considering levels of protein expression and reduction in endonuclease activity we chose the D693A mutant CCHFV-L for further structural and functional studies. Inhibition of endonuclease activity is important as downstream experiments require the addition of synthetic RNA which must remain undegraded.

### Molecular architecture of the CCHFV-L core

To obtain structural information we prepared a sample of the full-length CCHFV-L. The resulting CCHFV-L structure exhibited a preferred orientation; however, through extensive and iterative classification and repicking we could recover minor populations of orthogonal particle views from the data (Supplementary figure 2a, b). The resulting reconstruction was resolved to a global resolution of 2.74 Å with regions of the map in the core of the L-protein extending to 2.4 Å (Supplementary figure 2c). A model was built and refined into the map using an *in silico* structure prediction as a starting point.

The model encompasses residues 928-3226 and covers the endonuclease linker and the RdRp core of CCHFV-L (Fig 2a-c). The region of the map likely corresponding to residues 1901-2263 appeared highly mobile. Although it could be observed at a low map threshold, the quality was insufficient to build this region of the model, despite extensive classification or flexible refinement. N-terminal regions containing the OTU and endonuclease domains or C-terminal domains were not observed.

**Figure 2.**
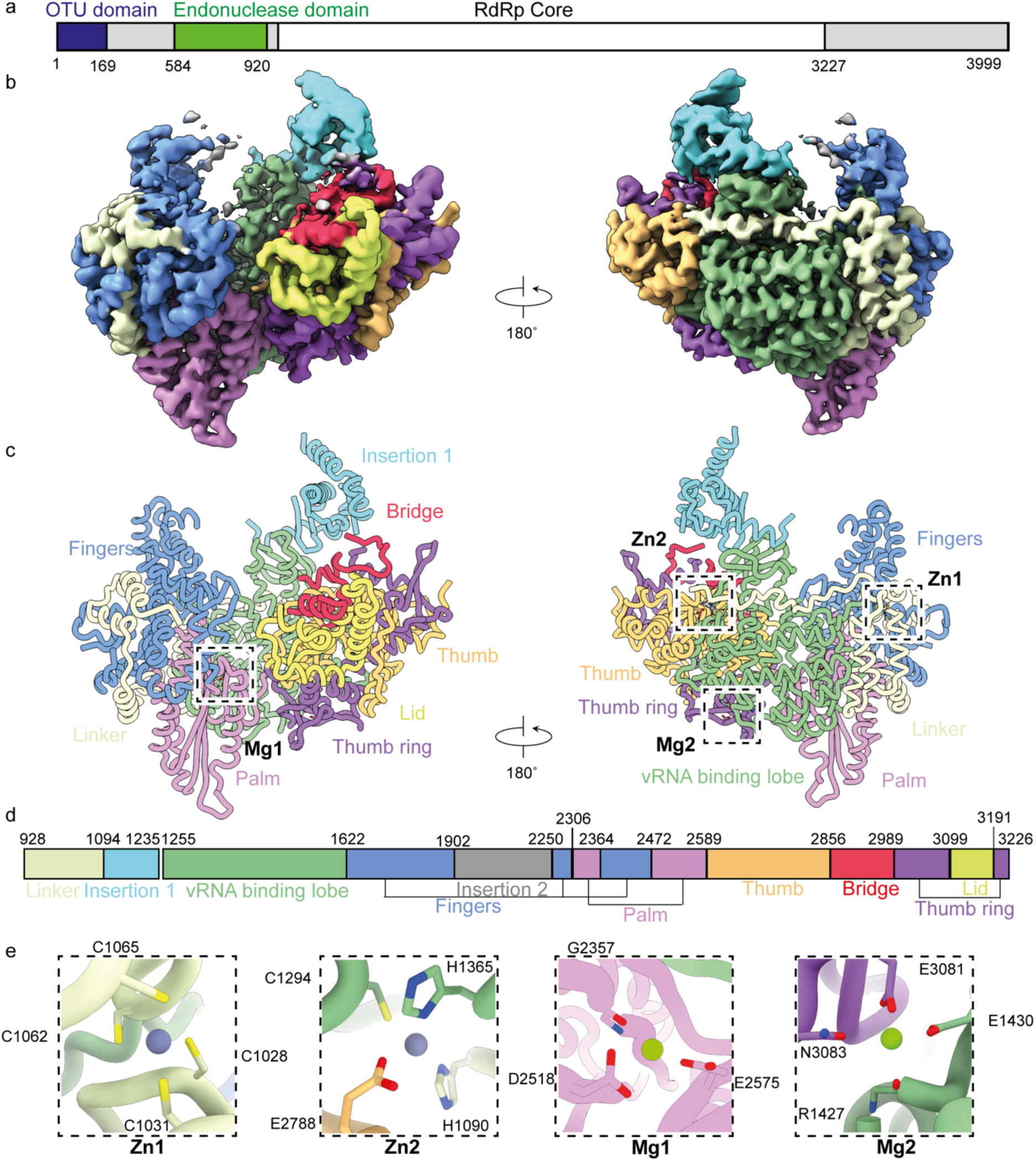
Molecular architecture of the CCHFV-L core. a) Domain arrangement of the full-length CCHFV-L. b) Density map and c) molecular model of the CCHFV-L RdRp. Sites of metal coordination are highlighted. d) Domain boundaries and annotation for the structure presented in the above panels. e) Metal coordination sites are shown in zoom with interacting residues annotated.

From the N-terminus of our model we observe a 170 residue long endonuclease linker (residues 928-1093) that winds around approximately half of the RdRp (Fig 2c, d). The linear path of the linker is interrupted by a three-helix zinc finger domain coordinating zinc ion 1 (Fig 2e). This zinc ion is tetrahedrally coordinated by C1028/1031 at the N-terminus of one helix and C1062/1065 located at the C-terminus of the second helix. At the C-terminus of the linker a second zinc ion is coordinated by H1090, C1294 and H1365 from the vRNA binding lobe (VBL), and E2788 from the thumb domain (Fig 2e). One zinc finger domain has been previously reported in the first 1325 residues of CCHFV-L^30^ and our data suggest it is likely those coordinating zinc ion 1. The structural role of these ions appears to be important in tethering the endonuclease linker tightly to the core of the RdRp. While structural metal ions are uncommon in polymerases of segmented RNA viruses^13^, they are frequently observed in the L-proteins of non-segmented RNA viruses^31^. The residues involved in metal coordination also appear well conserved within nairoviruses (Supplementary figure 3).

Joining the endonuclease linker and the VBL is an additional folded domain, which we term insertion 1 (residues 1094-1235). A break in the density suggests that it is flexibly associated with these domains. Insertion 1 is formed by five α-helices and a three stranded β-sheet. We could not identify a structural homologue to the domain in other bunyavirus L-proteins. Insertion 1 occupies a similar position to that of the pyramid domain from Arenavirus L-proteins^13^, suggesting it may perform a comparable yet to be determined role. Similarly, this could be an expanded clamp region which has been observed in several bunyavirus L-proteins^17,18^. In Tomato Spotted Wilt Virus L-protein the clamp region comprises approximately 50 residues and undergoes an ordered/disordered rearrangement upon RNA binding^18^. We have not observed a similar transition in our models; however, they do not contain the 3’ genomic RNA which may be important for this rearrangement.

Residues 1255-1621 form the VBL that constitutes the binding site for the 5’ end of the viral genome. Residues 1235-1256 which link insertion 1 and the VBL are disordered. The VBL is followed by the tripartite fingers domain (residues 1622-1901, 2250-2306, 2365-2472) which is concatenated with the bipartite palm domain (residues 2307-2364, 2473-2589) (Fig 2c, d). In many bunyavirus L-proteins both the fingers and the palm domains are bipartite, however, CCHFV-L has an extra insertion between residues 1902-2249 which was poorly resolved in the map. We have termed this insertion 2, and while it is not ordered in our maps, low threshold density suggests that it is flexibly located above the RdRp active site. Structural searches revealed no detectable homology to enzymatic domains, suggesting a structural role.

Annotation of the RdRp core highlights the location of motifs A-G which are collectively critical for RNA and NTP binding (Supplementary figure 4a, b). Motif F, likely at residues 2272-2298, was only partially resolved. The RdRp active site contains the canonical double aspartate residues responsible for active site metal coordination, with residues 2517 and 2518 (Supplementary figure 4b). We observed a magnesium ion in the RdRp active site tightly coordinated by residues D2518, E2575, and the backbone carboxyl of G2357 (Fig 2e). The Motif I, centred on residue K1620, is found in the hinge region between the fingers and VBL domains where it is predicted to interact with motif F.

Opposing the fingers domain are the thumb (residues 2589-2856), thumb ring (residues 2989-3099, 3191-3226), and lid (residues 3100-3190) domains. Another metal ion is coordinated by two residues from each the VBL and the thumb ring domain (Fig 2e). The coordination by these four residues suggests an octahedral geometry, consistent with magnesium as the most likely metal. Residues 2857-2989 contains the bridge domain that links the thumb and thumb ring domains. In our model we observe three stretches of ordered residues, however approximately 60% of the residues in this region are not ordered. This region potentially contains the template exit loop and the priming loop.

### Binding of the 5’ genome to the CCHFV-L core

We next sought to characterise the binding of the end of the 5’ genome to CCHFV-L. Purified CCHFV-L was mixed with a 15 times molar excess of 16mer synthetic RNA from the 5’ terminus (5’-UCU CAA AGA UAU AGC A-3’) and incubated on ice for 5 minutes. Immediately prior to preparing cryoEM grids a four times molar concentration of nanobody Nb20096 was added to help prevent potential issues with preferential orientation. From this dataset we were able to determine a complex describing the binding of the 5’ genome terminus, the CCHFV-L core, and the Nb20096 binding site to a global resolution of 2.31 Å (Fig 3 a, Supplementary figure 5a-c).

**Figure 3.**
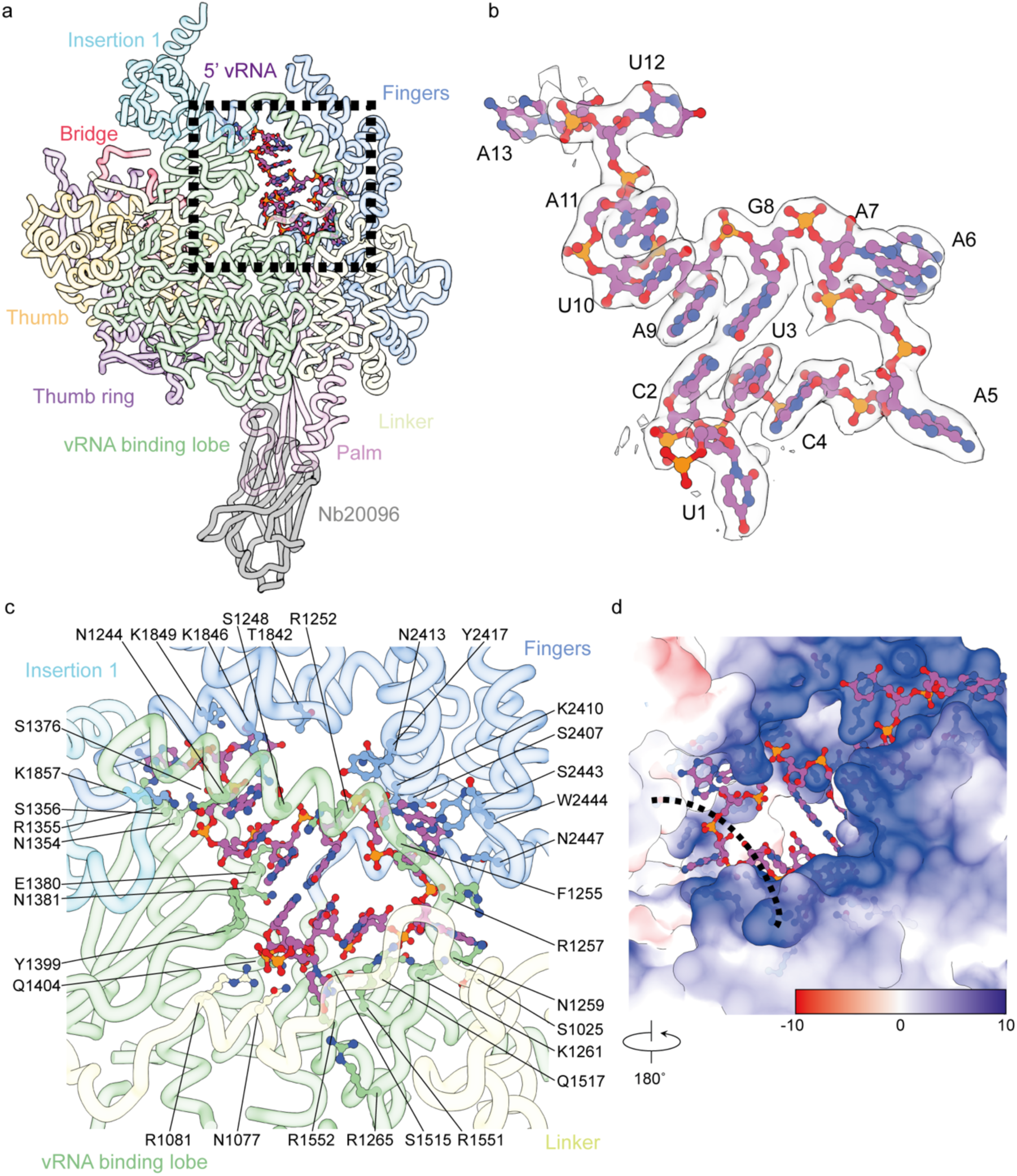
Coordination of the 5’ genome termini within the CCHFV-L. a) The molecular model of the CCHFV-L to the 5’ genome termini in complex with Nb20096. b) Density map of the 5’ vRNA with the model fit (purple). Base numbers and identity are annotated. c) Detailed molecular model showing the residues important in coordinating the RNA. Residues are shown in sticks and coloured according to the respective CCHFV-L domain colour code. d) Electrostatic potential surface representation of the RNA binding site highlighting strong electropositivity. The map is shown as a 180°rotation of panel c. The dashed line shows where residues have been removed to show the charge of the RNA binding site with more clarity.

The 13 bases at the 5’ terminus of the genome were observed to occupy a site formed by the endonuclease linker, fingers domain, VBL, and insertion 1 domain (Fig 3a-c). The RNA-protein interface buries an interface of approximately 2100 Å^2^. The 11 bases at the 5’ terminus form a tight hook structure, stabilised by a single Watson-Crick base pair between the cytosine at position 2 and the guanosine at position 8. Bases 2-4 form a three-base stack with the base in position 4 also forming a π-stacking interaction with the head group of R1551. The adenosine nucleotides in positions 5-7 have their bases flipped out with each base accommodated into a specific pocket in the CCHFV-L (Fig 3c). Bases 8-11 form a four base stack that is stabilised by large regions of the VBL and fingers domain. Nucleotides 12 and 13 are outside of the hook structure and less well coordinated into CCHFV-L, making fewer and less base specific interactions. The final three bases present in the RNA were not ordered likely due to limited interactions with the CCHFV-L. Analysis of the electrostatic surface of the hook-binding site reveals a strongly electropositive region, as expected for coordinating the negatively charged RNA backbone (Fig 3d).

Nb20096 binds to the palm domain, burying approximately 812 Å^2^ of surface area. The interface is maintained by residues from each of the three complementarity-determining regions (CDR) (Supplementary figure 4c, d). Notably, we observed a non-canonical interaction involving the residues N-terminal to CDR3, outside of the highly variable region, which contribute to a large area of the interface with the palm domain.

Comparison of the 5’ vRNA bound and RNA free CCHFV-L models allow us to understand the changes caused by RNA binding. The 5’ promoter RNA is stabilised through the ordering of residues 1236-1256 in the VBL. These residues function similarly to the arch from the PA subunit of influenza virus polymerase^32^, Tomato Spotted Wilt Virus^18^ and Hantavirus L-proteins^20^. In addition to the residues 1843-1859, 1870-1896 from the fingers domain have also become ordered and pack on top of the arch and directly onto the RNA hook. Residues at the hinge region between the VBL and the fingers domain (residues 1622-1628) have also become well ordered, suggesting a rigidification of the CCHFV-L core (Fig 4a). In neither of the 5’ promoter bound or RNA free models do we observe the full motif F structure, whereas previous studies have shown RNA binding to cause a rearrangement of this motif and an activation of the RdRp activity^17,18,22^. Similarly, neither model contained an ordered priming loop. Interestingly, previous activity studies of CCHFV-L have shown that addition of the 3’ template alone was a suitable template for efficient extension^33^, suggesting the 5’ promoter may not have the activating effect seen for other bunyaviruses.

**Figure 4.**
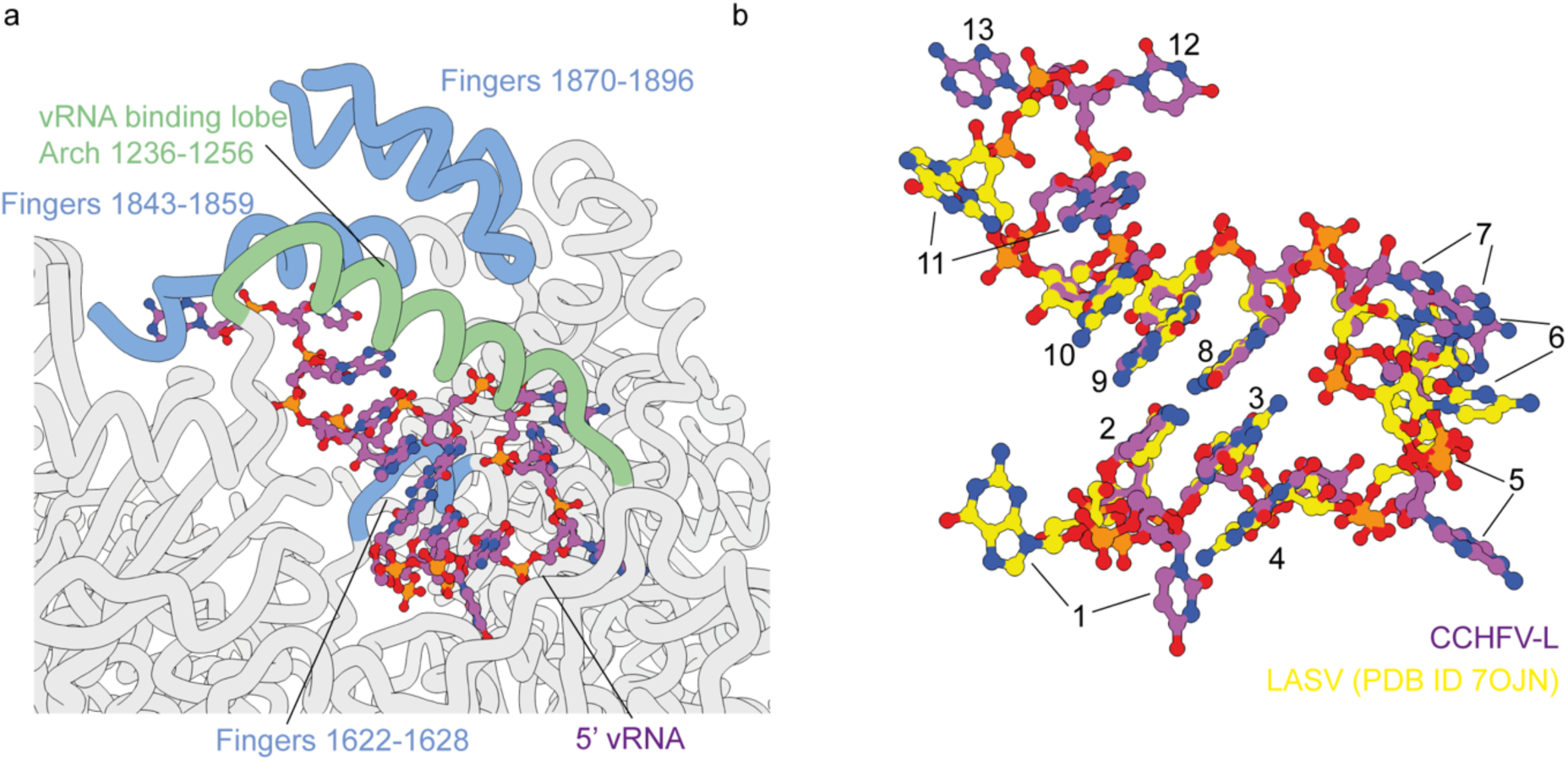
Effect of RNA binding and the 5’ hook structure. a) Comparison of the Apo and 5’ RNA bound structures. Regions which are ordered upon RNA binding are highlighted for the fingers (blue) or vRNA binding lobe (green). RNA is shown (purple). b) Structural comparison of the 5’ RNA terminus between CCHFV (purple) and LASV (yellow) show strong similarity with conservation at bases 2-4 and 8-10 maintaining base pairing and stacking interactions.

Comparison of the 5’ promoter structure to that of other segmented RNA viruses shows the strongest homology to Lassa virus^13^. Similarities include the first base of the genome protruding out from the L-protein, the hook structure being maintained by a single C-G base pair between positions 2 and 8, and nucleotides 5-7 having extensive base-protein contacts between (Fig b). In contrast, other bunyavirus 5’ promoters have between 2-4 base pairs that stabilise the hook structure.

### Initiation of viral genome replication

We next sought to assess the activity of the RdRp within the full-length CCHFV-L protein. For this purpose, we developed a ^32^P-labeled radionucleotide extension assay. Previous work has shown that the polymerase efficiently extends an RNA mimic of the 3’ template^29^. As our structural analysis demonstrated that the 5’ terminus forms an integral component of the CCHFV-L core, we included both 5’ and 3’ termini in our assay templates. These RNAs contained the 13 bases from the 5’ terminus and the 15 bases from the 3’ terminus of the genome. Previous analyses have shown that within the 5’ promoter, the nine terminal bases are highly conserved, as is a stretch from bases 17-21^34^. To mimic the interaction between the 5’ and 3’ RNA segments, we introduced a 5’-GCGCGC… sequence to the 3’ RNA template and the complement to the 5’ RNA template, producing a 21 nt long template (Fig 5a).

**Figure 5.**
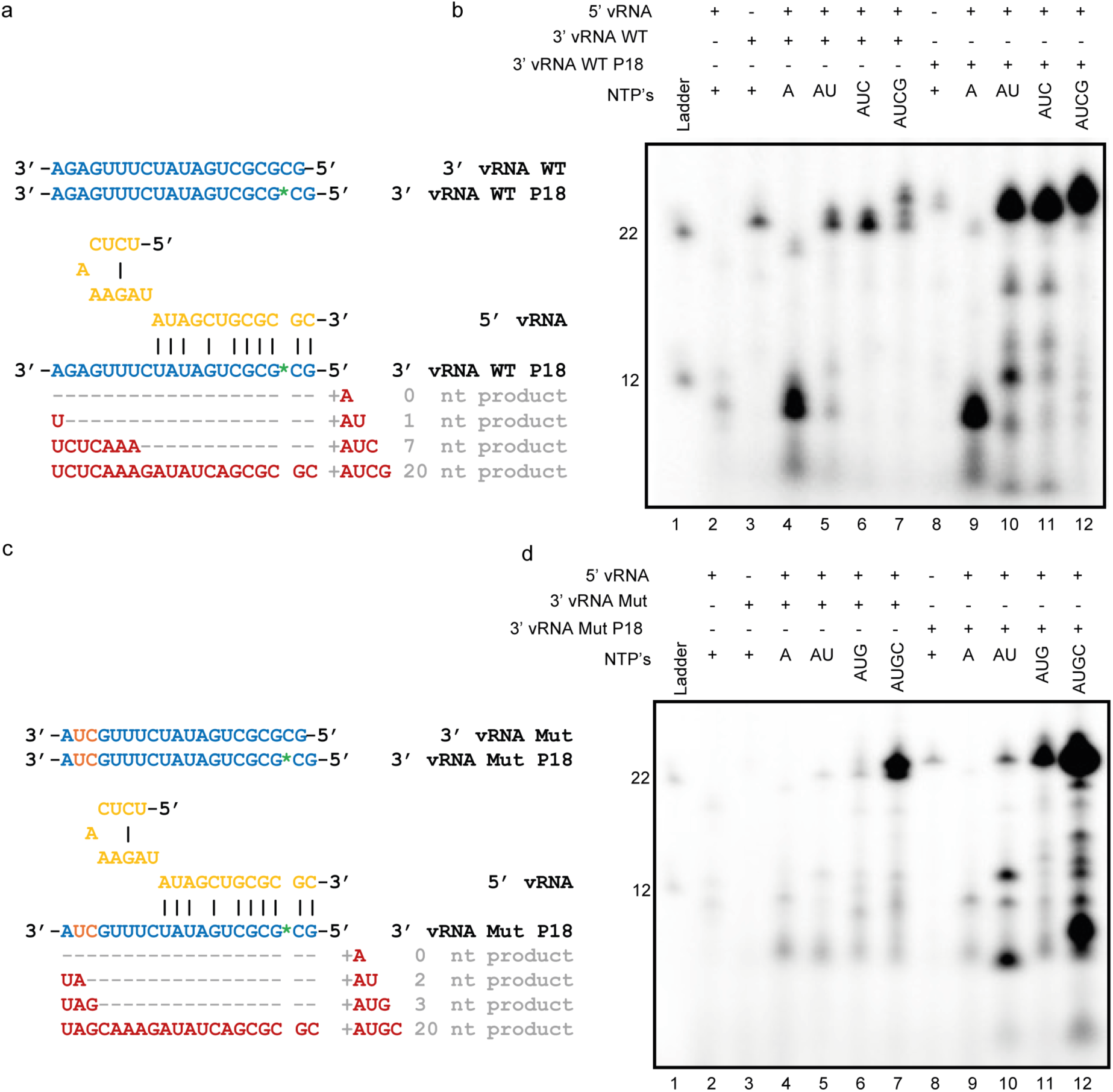
Unprimed radiolabelled extension assays. RNA templates and expected products yielded in the presence of different NTP combinations are shown (a) and (c). Location of the Cy3 fluorophore is indicated as a green asterisk. Reaction products generated from b) wild-type (WT) RNA or d) mutant (Mut) RNA templates. Base exchanges in in the mutant RNA template are highlighted in orange. Reaction conditions are detailed in the methods. Assays were performed in duplicate with comparable results.

Using wild-type templates corresponding to the 5’ and 3’ termini of the genome (Fig 5a), we observed that only limited extension products were produced (Fig 5b, lanes 2-3). Addition of ATP yielded an abundant ∼10 nt long product, with the further addition of UTP resulting in a product of approximately 22 nt, close to the expected full-length size (Fig 5b, lanes 4-5). Further addition of CTP or GTP caused only minor changes in length and abundance of the observed products (Fig 5b, lanes 6-7). The observation of a 22 nt product in presence of only ATP and UTP suggests low fidelity of CCHFV-L under our experimental conditions.

Using a mutant 3’ promoter (3’ vRNA Mut) where bases 2 and 3 are mutated from GA to UC (Fig 5c) allowed us to control early stages of *de novo* initiation more precisely. In this system, only RNA products shorter that 3 nt would be produced unless all four NTP are present. Assays in the presence of a limited set of NTPs exhibited only low product abundance (Fig 5d, lanes 4-6), while addition of all NTPs promoted generation of the ∼22nt product (Fig 5d, lane 7). Assays performed with wild-type or D693A mutant CCHFV-L showed similar activities in these assays while CCHFV-L, whilst D2517A/D2518A active site mutations inhibited all activity (Supplementary figure 6c).

During assay development, we tested the activity of CCHFV-L on template RNAs containing fluorophores at several positions (Supplementary figure 6a). Incorporation of a Cy3 dye at base 18 of the wild-type 3’ template (3’ WT P18 vRNA) substantially enhanced product formation for both full-length and shorter products generated by the addition of ATP and UTP, or ATP, UTP and CTP (Figure 5b). Fluorophores introduced at other positions inhibited or reduced activity of CCHFV-L (Supplementary figure 6b). In reactions using mutated 3’ templates with Cy3 incorporation at position 18 (3’ Mut P18 vRNA), the addition of ATP and UTP strengthened the production of a ∼12 nt RNA product, whilst addition of the complete NTP set resulted in a ladder-like production of shorter transcripts alongside the strong full-length transcript (Figure 5d, lanes 10 and 12). We speculate that the increase in activity observed due to incorporation of the dye at position 18 is related to a destabilisation of the predicted RNA duplex, perhaps enabling optimal positioning of the template RNA.

### Development of a primed fluorescent extension assay

Our unprimed assays suggested strong polymerase activity; however, we suspected that the early stages of initiation may be prone to non-templated extension. To prevent initiation, we developed a primer-dependent assay using a 5 nt RNA primer (Fig 6a). We synthesised RNA primers without a label and with a Cy5 fluorophore attached to the 5’ of the U1 nucleotide (P1-Cy5 primer).

**Figure 6.**
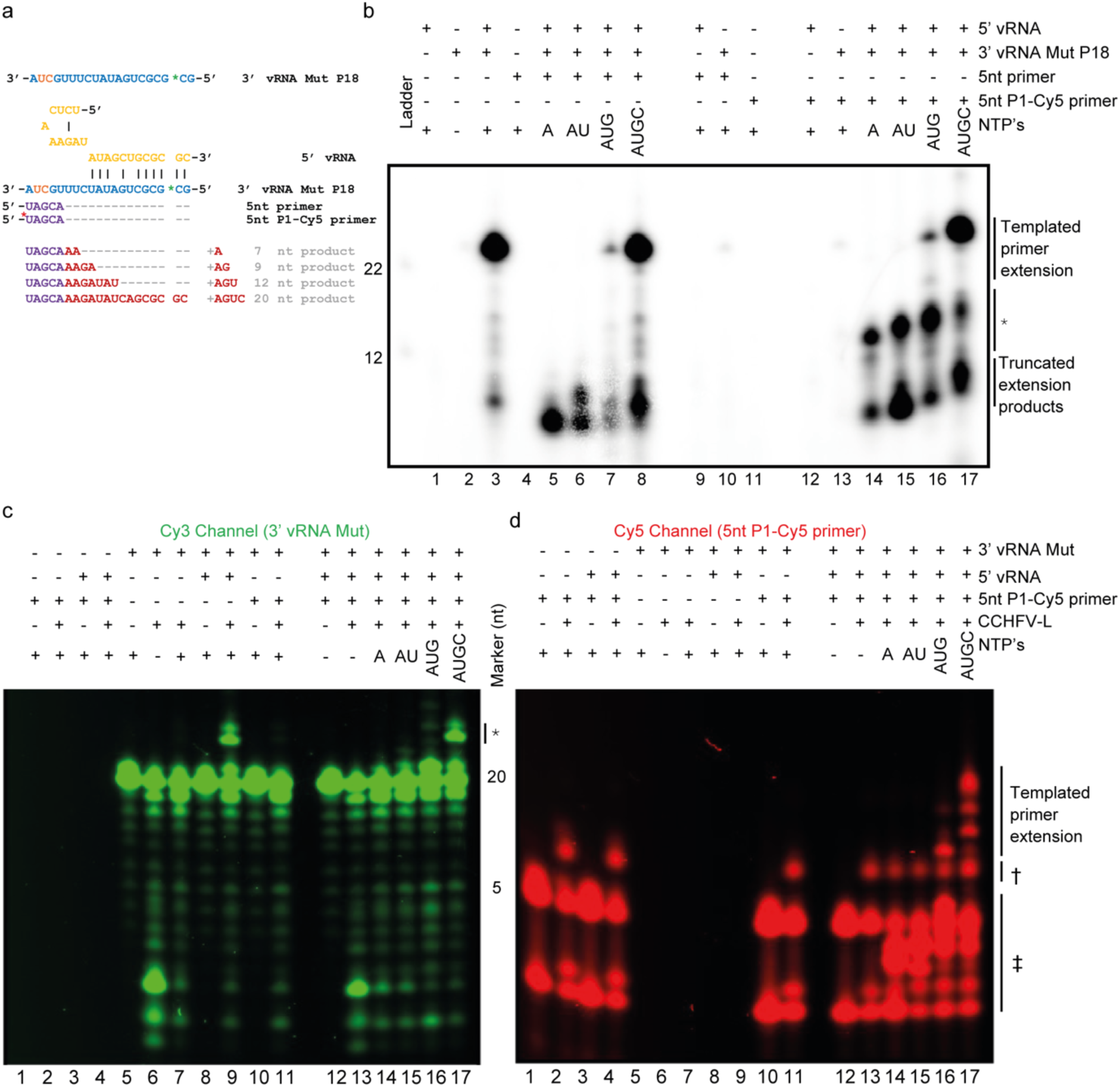
Primed radiolabelled and fluorescent extension assays. a) RNA templates and expected products produced in the presence of NTP combinations with the addition of a 5 nt RNA primer. Location of fluorophores are indicated with a green (Cy3) or red (Cy5) asterisk. b) Reaction products generated from a 3’ vRNA Mut P18 and 5’ vRNA templates with either the unlabelled 5 nt primer (lanes 4-10) or the 5 nt P1-Cy5 primer (lanes 11-18). * A >12nt reaction product appearing only in the presence of 3’ vRNA, 5’ vRNA, and P1-Cy5 primer is likely due to partial extension of the primer with ATP alone. c-d) Fluorescent reaction products generated from the Mut P18 vRNA template together with the 5’ vRNA template and P1-Cy5 primer, split into Cy3 and Cy5 fluorescence channels. *A Cy3-labelled fluorescent extension product in the presence of Mut P18 vRNA, CCHFV-L and 5’ vRNA likely indicates non-templated extension of the 3’ Mut P18 vRNA. † A Cy5-labelled band over 5 nt in length is the product of CCHFV-L and P1-Cy5 primer in the absence of NTPs, indicating an interaction between the L-protein and P1-Cy5 primer independently of RdRp activity. ‡ Cy5-labelled fluorescent bands shorter than 5 nt are likely derived from extension using a degraded 5 nt P1-Cy5 primer. Assays were performed in duplicates with comparable results.

Reactions including only the 3’ template and primer resulted in very reduced product formation (Fig 6b, lanes 10 and 13). Further inclusion of the 5’ strand enhanced the production of the 22 nt product (Fig 6b, lanes 8 and 17), suggesting a strong activating role for the 5’ RNA template. Previous studies have shown extension using just a 5 nt primer and 3’ template RNA^29^, however our structural data suggests that the 5’ end is also required, a concept which is strongly supported by our functional data.

Withholding nucleotides in these assays limited the extension of products when reactions were initiated with the unlabelled primer (Fig 6b, lanes 5-8). On addition of ATP we observed a dominant short reaction product at ∼7 nt in length. We noted another stronger extension product at ∼9 nt on further addition of UTP, whilst further addition of GTP produced a weak full-length reaction product. Using all NTPs, we observed a strong full-length product with additional short extension products being visible.

To differentiate between de novo extension products and primed extension products, we developed a fluorescence extension assay using the P1-Cy5 primer. Upon sequential addition of NTPs, we observed the production of increasingly longer products similar to the reactions using unlabelled 5 nt primer (Fig 6b, lanes 14-17). In addition, we observed an abundant intermediate product at the length of approximately 12 nt. We believe this product is directly related to the fluorescent primer, as the presence of the fluorophore is the only difference between the reactions. The molecular identity of the product is unclear, though it may be related to synthesis issues producing a degraded product.

We next replicated the conditions of the radiolabelled extension assay, however using the fluorescently labelled 5 nt primer to observe RNA products (Fig 6c, d). We could clearly observe the extension of the primer in ATP/UTP/GTP conditions, producing an extension product of ∼13 nt. Addition of CTP produced a strong fluorescent product at ∼20 nt in length, indicating a full-length product accompanied by intermediate products.

In combination the ^32^P radiolabelled and fluorescence extension assays show that the 5’ genome terminus potently enhances RdRp activity compared to single 3’ vRNA template-initiated products. We have also demonstrated that CCHFV-L could incorporate non-templated nucleotides into the extending product, and that CCHFV-L can directly add nucleotides to template RNA.

### Screening of nucleotide triphosphate analogues

We next turned our attention to inhibiting the RdRp domain of CCHFV-L using commercial nucleotide analogues. Ribavirin-TP and Favipiravir-TP were selected as broad-spectrum purine nucleotide analogue antivirals with demonstrated activity against RNA virus polymerases^35,36^. Each compound was screened for activity in our ^32^P radiolabelled extension assay at a final concentration of 1.25 mM (Fig 7a, lanes 4 and 5). Addition of Ribavirin-TP and Favipiravir-TP at high concentrations did not reduce the full-length product. However, Favipiravir-TP addition enhanced the formation of a truncated product at less than 12 nt in length. The titration of lower compound concentrations had no effect (Supplementary figure 6d). This activity is in line with the known modes of action of both Ribavirin and Favipiravir, which are believed to induce stalling rather than chain termination after their incorporation into the viral genome^35,37^.

**Figure 7.**
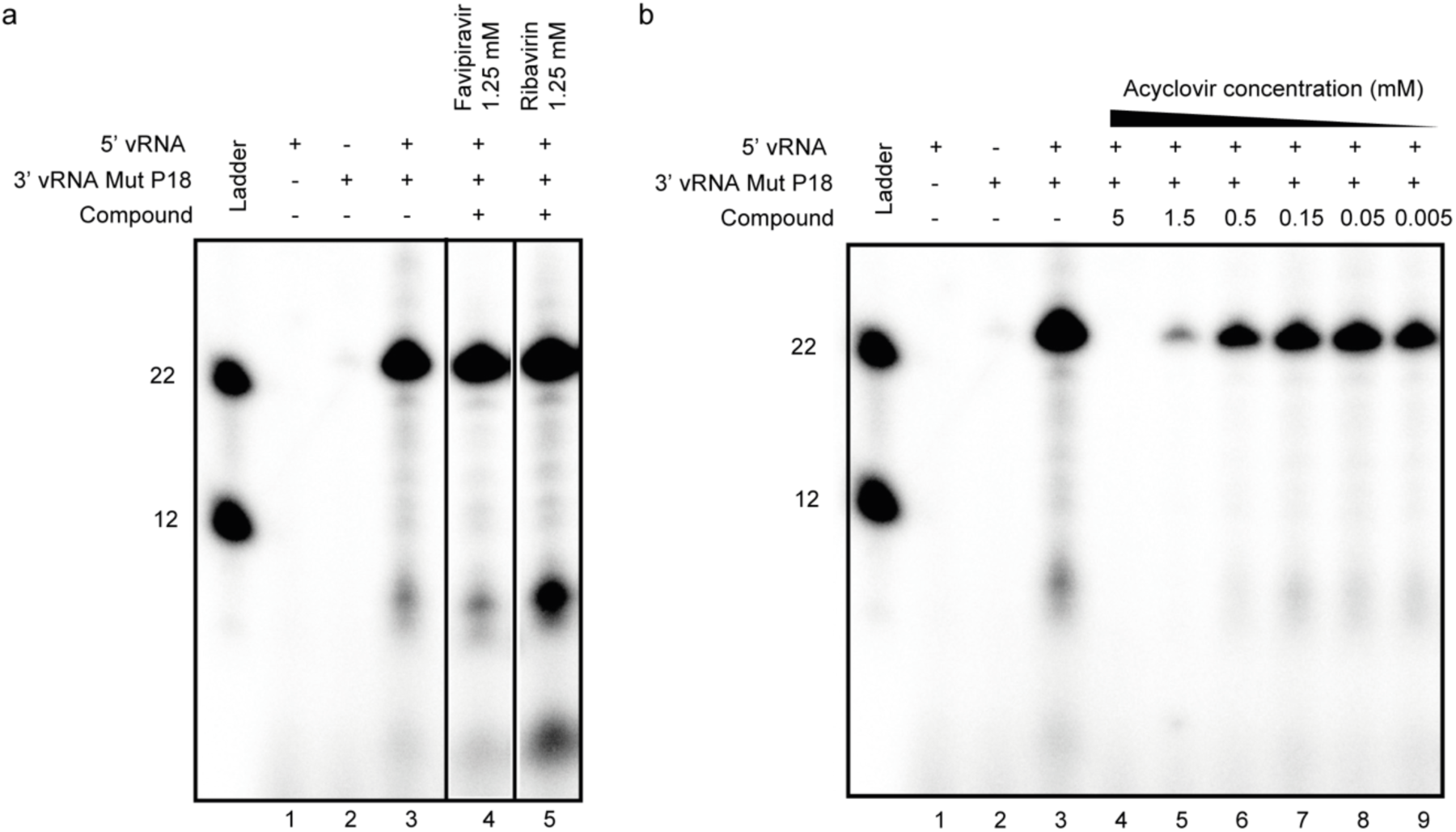
Screening of antiviral nucleotide triphosphate analogues. a) Reaction products generated from 3’ vRNA Mut P18 and 5’ vRNA templates in the presence of Ribavirin-TP and Favipiravir-TP at 1.25 mM concentration. The assay was run twice in parallel on the gel, producing the same result. b) Reaction products generated from 3’ vRNA Mut P18 and 5’ vRNA templates in the presence of Acyclovir-TP. Acyclovir-TP concentration was titrated from 5 mM to 5 µM. All lanes contain DMSO at 25% v/v. Reaction conditions are detailed in the methods. Assays were performed four times with comparable results

We next screened Acyclovir-TP, a chain terminating acyclic purine nucleoside analogue used to specifically target the DNA polymerase of herpes viruses since the 1980’s^38^. We titrated Acyclovir-TP into the radionucleotide extension assays from 5 mM to 5 μM concentrations (Fig 7b) showed complete inhibition of full-length product formation at the highest concentration (Fig 7b, lane 4), while at lower concentrations we only noted a partial reduction.

## Discussion/Conclusions

All bunyavirus L-proteins are thought to contain certain key features including an endonuclease domain, an RdRp domain, and a cap-binding domain. For most bunyaviruses with animal hosts the resulting L-protein is approximately 250 kDa. Here we have provided the characterisation of the function for two of the enzymatic domains and two high resolution structures describing the core RdRp of CCHFV-L.

The L-protein core we describe here is approximately 2150 residues which is much larger than the RdRp from structurally characterised bunyaviruses including arenaviruses (∼1550 residues^14^), hantaviruses (∼1340 residues^21^), peribunyaviruses (1470^16^), or phenuivirises (1340 residues^11^) (Supplementary figure 7). This extra mass is largely accounted for by two unexpected structural domains that we have termed insertion 1 and insertion 2, which are approximately 140 and 350 residues, respectively. Comparison to structures of other bunyavirus RdRps shows that the TSWV-L^18^ and LASV-L^14^ each have insertions into the RdRp, in a similar location as the insertion 1 in CCHFV-L (Supplementary figure 7). In TSWV-L the clamp domain is ordered upon addition of vRNA, while in our models it is similarly arranged in the presence or absence of RNA. We observe no structural homology between the insertion 1 and clamp domains, it appears only the insertion into the RdRp is conserved. The Insertion 2 domain in CCHFV-L appears conserved within nairoviruses (Supplementary figure 3) but is not apparent in other bunyaviruses. In our models it is poorly ordered, and we are unable to determine its functional role. Comparison of the endonuclease linker from other bunyavirus L-proteins demonstrates that CCHFV-L is distinct in containing a zinc finger domain (Supplementary figure 7), the functional importance of this is yet to be understood.

Our functional data reveal that the activity of CCHFV-L is potently activated by the addition of the 5’ genome RNA, generating a strong increase in the replication activity of the protein. The more distantly related influenza virus polymerase has been demonstrated to show a strong increase in activity with the addition of promoter RNA^39^. Studies on the more closely related arenavirus L-proteins showed that the addition of the 5’ vRNA promoter increased the activity of the LASV-L in de novo replication but reduced activity in cap-primed assays^14^. Given the comparable promoter structure of Arenaviruses and CCHFV-L (Fig 4 b), it may be that they function in a similar manner in CCHFV.

Our antiviral screening experiments unexpectedly identified Acyclovir-TP as an inhibitor of the CCHFV-L (Fig 7b). Typically, Acyclovir is used against Herpes viruses where in the triphosphorylated form it acts a potent chain terminator, inhibiting the function of the viral polymerase. We are yet to observe active production of RNA with cryoEM of the CCHFV-L, but future studies will aim to reveal the molecular details of this inhibition. Potentially this will open up novel approaches for antiviral design.

Collectively, we have provided a structural and functional description of a pathogenic nairovirus L-protein. These findings provide a fundamental increase in our understanding of CCHFV processes and provide new avenues for antiviral discovery.

## Methods

### Protein cloning and purification

The viral L-protein gene from *Orthonairovirus haemorrhagiae* (NCBI Reference Sequence: YP_325663.1) was synthesised (SynBio) with N-terminal twin-strep and octa-His tags followed by a TEV protease cleavage site in a pFastBac vector. Genes encoding mutations were generated using standard cloning techniques. Baculoviruses for all viruses were generated according to standard protocols in sf9 insect cells maintained in serum-free Sf-900 media (Gibco). For large scale protein expression cells were infected with virus and left for 72 hours. After harvesting the cell pellet this was resuspended in a buffer containing 50 mM HEPES, pH 7.5, 500 mM NaCl, 0.05% (w/v) n-Octyl beta-D-thioglucopyranoside, 2 mM dithiothreitol, 10% (v/v) glycerol, one protease inhibitor tablet (Sigma), 5 mg RNAse, and 1.2 mL of BioLock (IBA). After 30 minutes cells were further lysed by sonication, clarified with centrifugation and the supernatant incubated with Strep-Tactin Superflow high capacity (IBA) resin for 2 hours. Resin was washed with 100 column volumes of buffer comprising 50 mM HEPES, pH 7.5, 500mM NaCl, 0.05% (w/v) n-Octyl beta-D-thioglucopyranoside, 2 mM dithiothreitol, and 5% (v/v) glycerol. Prior to elution a final wash with 20 column volumes of 50 mM HEPES, pH 7.5, 500mM NaCl, 1 mM dithiothreitol, and 5% (v/v) glycerol. CCHFV-L was then eluted with 10 column volumes 20 mM HEPES, pH7.6, 500 mM NaCl, 1 mM dithiothreitol, 5% (v/v) glycerol, and 50 mM Biotin. As required CCHFV-L was passed over a Superose 6 Increase 10/300 column into a buffer containing 20 mM HEPES, pH7.6, 500 mM NaCl, 1 mM dithiothreitol, 5% (v/v) glycerol. The sample was then concentrated and snap frozen in liquid nitrogen prior to storage at -70°C.

### Nanobody generation and purification

Nanobody were generated against CCHFV-L as previously described^40^. Briefly, a plasmid containing the nanobody-20096 (Nb20096) gene with a C-terminal His tag, was transformed into chemically competent Escherichia coli WK6 cells. Cells were then grown in LB media supplemented with 0.1% glucose (w/v), 1 mM MgCl_2_, and 100 mg/mL ampicillin to a optical density of 0.7 prior to overnight induction with 1 mM isopropylthiogalactoside at 28°C. After centrifugation to collect the cells, nanobodies were released from the periplasm by osmotic shock. The supernatant was then clarified by centrifugation, bound to NiNTA resin (Qiagen), before elution with 500 mM imidazole. Once concentrated the elution was then applied to a Superdex S75 Increase 10/300 GL column equilibrated in 20 mM HEPES, pH7.6, and 150 mM NaCl. Fractions which contained the Nb20096 were again concentrated and stored at -20°C.

### Mass Photometry

Experiments were performed on a TwoMP mass photometer (Refeyn). Prior to conducting experiments, a calibration with three standards (thyroglobulin, ovalbumin, and aldolase) were each diluted in buffer containing 20 mM, HEPES, pH 7.5, 500 mM NaCl, 0.5 mM dithiothreitol, and 5% (v/v) glycerol. For CCHFV-L analysis the protein was diluted to a final concentration of 0.01 mg/ml in the same buffer. Analysis was performed using the DiscoverMP software.

### Endonuclease cleavages assays

A Fluorescein dye labelled 25-mer ssRNA substrate (5′-UAGUAGUAUGCUCCGCAGGAACAAA-3′), was chemically synthesised (Integrated DNA Technologies). Endonuclease activity assays were performed by incubating 2 μg CCHFV-L with 2.5 μM of 25-mer ssRNA and 2.5 mM of MgCl_2_. The reactions were incubated at 30°C at time intervals ranging from 0-180 minutes. At each time point, the sample tube was removed and reaction stopped by addition of RNA loading buffer to a final concentration of 45% formamide and 5 mM Ethylenediaminetetraacetic acid (EDTA) prior to heating samples to 95°C for 3 minutes. Reaction products were resolved by 7 M Urea 20% polyacrylamide Tris-borate-EDTA (TBE) gel electrophoresis (PAGE) in 0.5X TBE buffer. Fluorescent signal was detected using Chemidoc MP (Biorad). Uncropped images are provided in the source data file.

### Cryo-EM sample preparation

Building from our early work demonstrating the CCHFV-L contained an active endonuclease domain, our cryoEM experiments were prepared using the D693A mutant. All grids were prepared using a Vitrobot mark IV (FEI) at 100% humidity. UltrAuFoil (Quantifoil) (R2/2 on a 200 mesh or R1.2/1.3 300 mesh) grids were glow discharged, before a volume of 3.5 μL sample and blotted for 5.5 seconds before vitrification in liquid ethane.

The RNA free CCHFV-L structure was prepared by directly diluting frozen protein stock to a final concentration of 0.8 mg/mL with 20 mM HEPES, pH7.6, 500 mM NaCl, and 5% (v/v) glycerol.

The CCHFV-L-Nb20096-5’RNA sample contained CCHFV-L at a final concentration of 0.6 mg/mL (∼1.3 μM), Nb20096 at a concentration of 0.06 mg/mL (∼5 μM), and 20 μM 5’ RNA (5’-UCU CAA AGA UAU AGC A-3’). RNA was incubated with CCHFV-L for 5 minutes prior to the addition of Nb20096 and the grids prepared immediately.

### Cryo-EM image collection

Cryo-EM data for the RNA free CCHFV-L were collected at the Oxford Particle Imaging Centre, on a 300 kV G3i Titan Krios microscope (Thermo Fisher Scientific) equipped with a SelectrisX energy filter and Falcon IV direct electron detector. Data were collected automatically in EPU 3.4, movies were recorded in EER format with a total dose of ∼50 e-/Å^2^ and a calibrated pixel size of 0.932 Å/pix.

The CCHFV-L-Nb20096-5’RNA was collected at the EMBL Imaging Centre in Heidelberg, on a 300 kV G4 Titan Krios microscope (Thermo Fisher Scientific) equipped with a SelectrisX energy filter and Falcon IV direct electron detector using SerialEM version 4.2.0beta^41^. Data were collected in EER format with a total dose of ∼50 e-/Å^2^ and a pixel size of 0.73 Å/pix. As preferential orientation had been observed in screening samples 7445 movies were collected at a 0° before the stage was tilted to 30° and a further 5784 movies collected.

### Cryo-EM data processing

Data were processed using cryoSPARC V4.7.1 using the same initial process^42^. EER format movies were fractionated in 40 frames without applying an up-sampling factor. Patch motion correction and patch CTF-estimation were performed with default setting, movies with poor statistics were removed. To generate a first round of candidate particles for processing blob picker was used followed by 2D classification. These initial templates were then used for template-based picking. These two particle sets were merged, duplicates removed, 2D classified, and the classes containing high resolution information used to train a Topaz model^43^. Particles from the different picking approaches were then pooled, duplicates removed, and a first round of ab-initio models generated and subsequently refined using heterogenous refinement. Particles which classified into the high-resolution heterogenous classes were then used to retrain a Topaz model and the process was iterated twice. The final high-resolution heterogenous class was then further refined using non-uniform refinement (with per particle CTF refinement) followed by a local refinement. This map was then considered the consensus refinement. For the RNA free dataset this was where processing stopped. For the RNA bound dataset we performed further 3D classification to remove particles which did not have RNA bound before a final round of local refinement.

### Structure determination and model refinement

To generate an initial protein model the AlphaFold 3 server^44^ was used. The map and model were imported into UCSF ChimeraX^45^, globally positioned in the map and large regions not observed in the model removed. Cycles of manual refinement and rebuilding in Coot^46^ and real space refinement in PHENIX^47^ were used to generate a final model. For the 5’ RNA model, the RNA was built de novo in coot. Final model geometry and map-to-model comparison was preformed using PHENIX Molprobity. Once the model had been built for the RNA bound complex this was used as a template for generating the RNA free model with cycles of refinement in coot and PHENIX. All map and model statistics are described in Supplementary Table 1. Structural analysis and figures were prepared using UCSF ChimeraX ^45^.

### In vitro activity assays

Wild-type and D693A CCHFV L proteins were used in radiolabelled and fluorescent extension assays. In the radiolabelled assay, CCHFV L protein at a final concentration of 1 μM was initially diluted in a reaction buffer containing 20 mM HEPES pH 7.5, 100 mM NaCl, 1 mM DTT, 10% glycerol, 5 mM MgCl_2_ and 5 mM MnCl_2_. The diluted L protein was added to a reaction mixture with final concentrations of 3’ template, 5’ template, and 5nt primer RNA at 250 nM, RNAsin Plus (Promega) at 1U/µl, GTP, CTP and UTP at 125 μM, ATP at 250 nM, and [α-32P]-ATP (Revvity) at 0.2 µl/reaction, added to a final volume of 3 μl. Reactions progressed at 30°C for 1 hour before an identical volume of Formamide dye (90% formamide, 10 mM EDTA pH 8) was added, and samples were heated to 80°C for 5 minutes. Samples were loaded onto 6 M urea, 20% polyacrylamide denaturing gel (dimensions 16.5 × 28 cm (w x h), C.B.S. Scientific) and run at 450 V for 4 hours to resolve RNA products. Radioactive extension signal was quantified through phosphorimaging screen exposure and subsequent scanning (Typhoon FLA 9500). Acyclovir-TP, Favipiravir-TP and Ribavirin-TP were kind gifts from the Fodor lab. Compounds were dissolved to a final concentration of four times the concentration used in the assay in 100% v/v DMSO before being added to the reaction at the final concentration.

Fluorescent RNAs except for the P1-labelled 5 nt primer (Dharmacon) were ordered from Integrated DNA technologies (IDT). Cy3, and Cy5 fluorophores were conjugated to the oligonucleotides through phosphoramidite chemistry for 5’, 3’, and internally positioned fluorophores.

The fluorescent extension assay proceeded as described for the radiolabelled extension assay, except that unlabelled ATP instead was added at a final concentration of 125 μM. Fluorescent signal was visualized using the Chemidoc imaging system (Bio-Rad), with separate Cy3 and Cy5 channels used to discriminate between the two fluorophores.

## Data availability

CryoEM maps and models generated in this study have been uploaded to the PDB and EMDB. The accession codes for the RNA Free model are 9T0F and EMD-55400. For the RNA bound model they are 9T0E and EMD-55399. Source data are provided as a source data file.

## Acknowledgements

We thank members of the Grimes lab for helpful comments and discussions and James Seaton for some preliminary modelling experiments. This work was supported by a Wellcome Investigator Award 200835/Z/16/Z (to J.M.G.). J.R.K was supported by a University of Oxford-Medical Sciences Division Pump Priming Award, a Royal Society Research Grant (RGS\R2\242315), and a Warwick Research Development Award with N.R. (RD24302). ISIDORe and INSTRUCT-ERIC supported access to the electron microscopes at EMBL Heidelberg (Project ID 26743). We acknowledge the support and the use of resources of Instruct-ERIC (PID24637, VID 42576), part of the European Strategy Forum on Research Infrastructures (ESFRI), and the Research Foundation - Flanders (FWO) for their support to the Nanobody discovery. We thank Eva Beke for the technical assistance during Nanobody discovery. R.C and N.R were supported by a Royal Society Dorothy Hodgkin Research Fellowship DKR00620 (to N.R). We thank Dr Simon Fromm from EMBL Heidelberg for assistance with data collection. Computational aspects of this work were enabled by the Oxford Biomedical Research Computing (BMRC; DOI-2025) facility. For the purpose of open access, the author has applied a Creative Commons Attribution (CC-BY) licence to any Author Accepted Manuscript version arising from this submission. Correspondence and requests for materials should be addressed to J.R.K.

## Author contributions

A.D, R.C, L.C, F.G., D.N.D, N.R, J.M.G, and J.R.K performed experiments and analysis. E.P and J.S provided nanobody materials. J.R.K wrote the paper with input from all authors.

---

**Supplementary Table 1.**
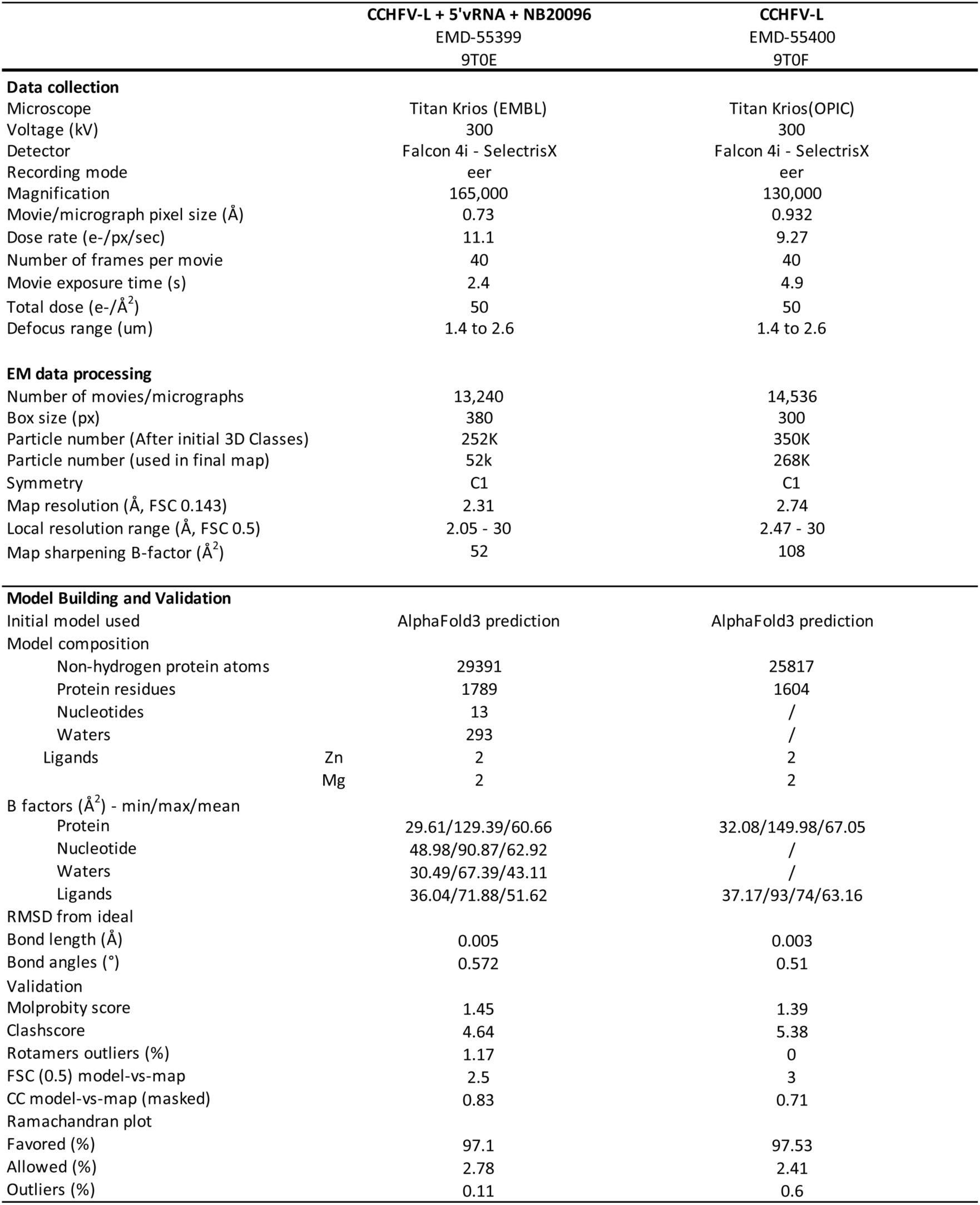
Cryo-EM collection and refinement parameters.

**Supplementary figure 1.**
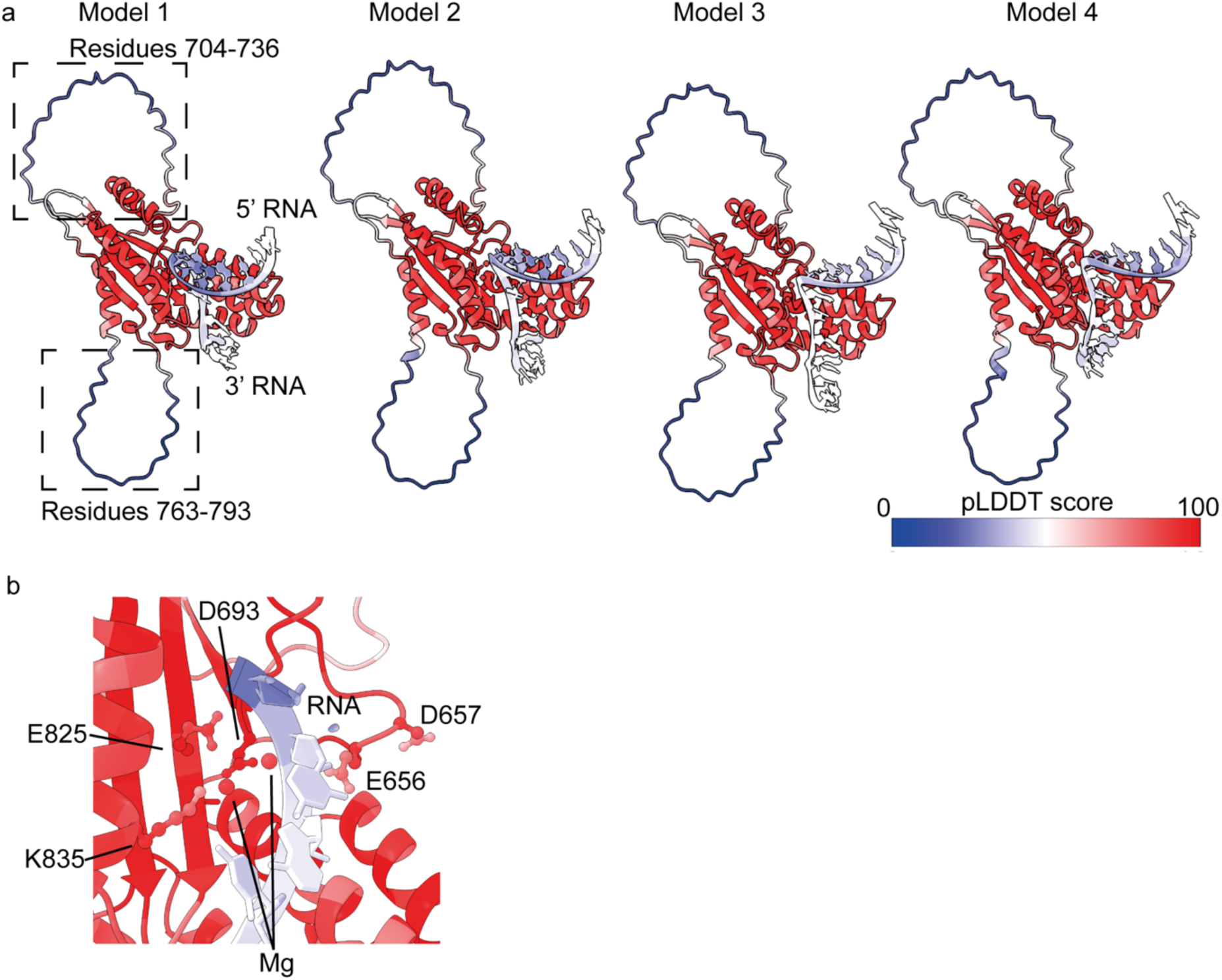
CCHFV endonuclease predictions bound to RNA and magnesium. a) Four models of the endonuclease domain have been predicted using the Alphafold Server. Models are coloured according to the pLDDT score. Residues of the endonuclease which are predicted with poor accuracy are shown in boxes. b) Zoom to the endonuclease domain active site showing RNA (sticks), magnesium (sphere), and potentially important residues (sticks). Colouring as per the pLDDT score shown in panel a).

**Supplementary figure 2.**
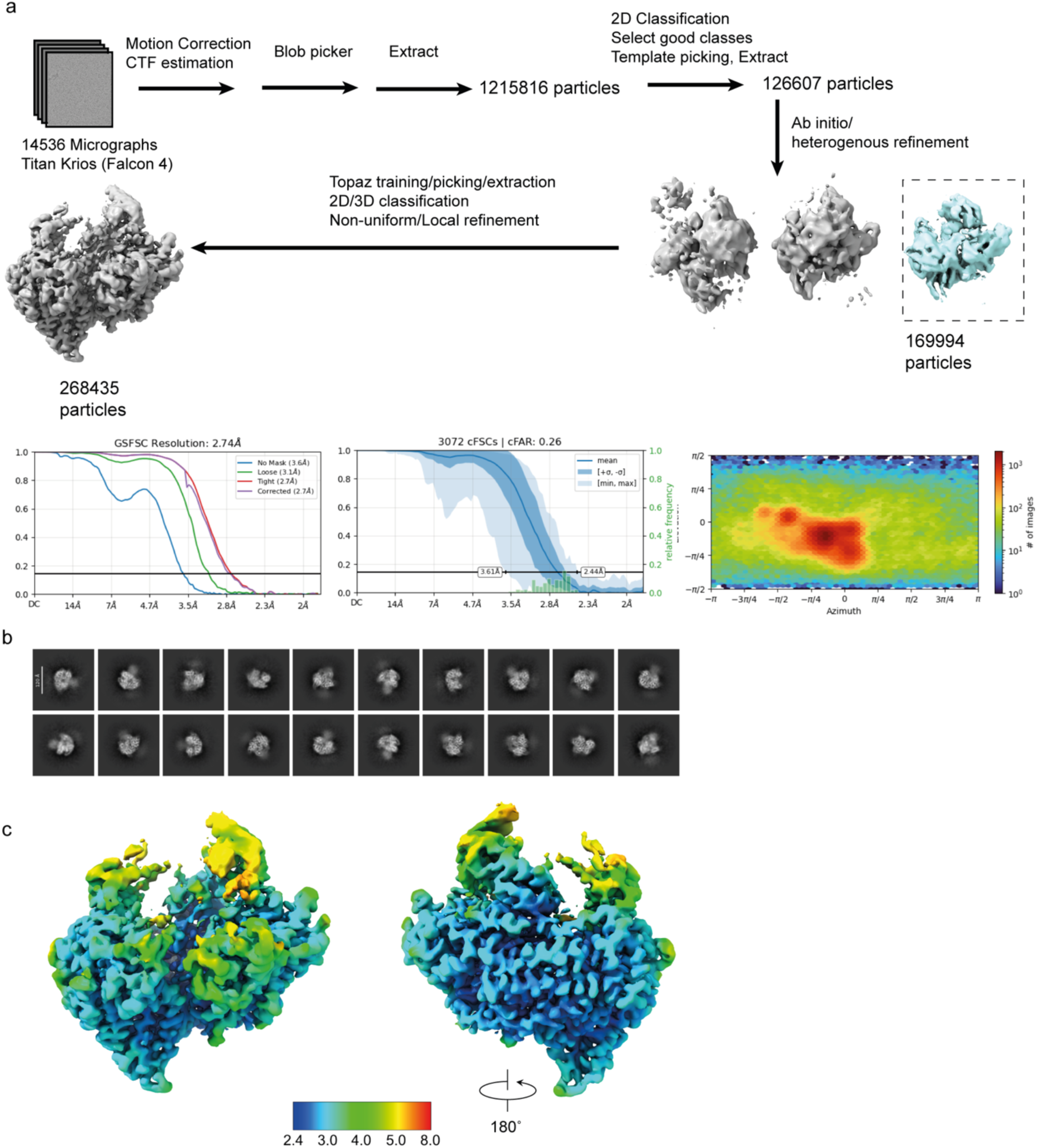
Cryo-EM processing workflow for RNA free CCHFV-L. a) Processing scheme for the RNA free CCHFV-L. b) 2D classes of CCHFV-L. c) Resolution range of the resulting map.

**Supplementary figure 3.**
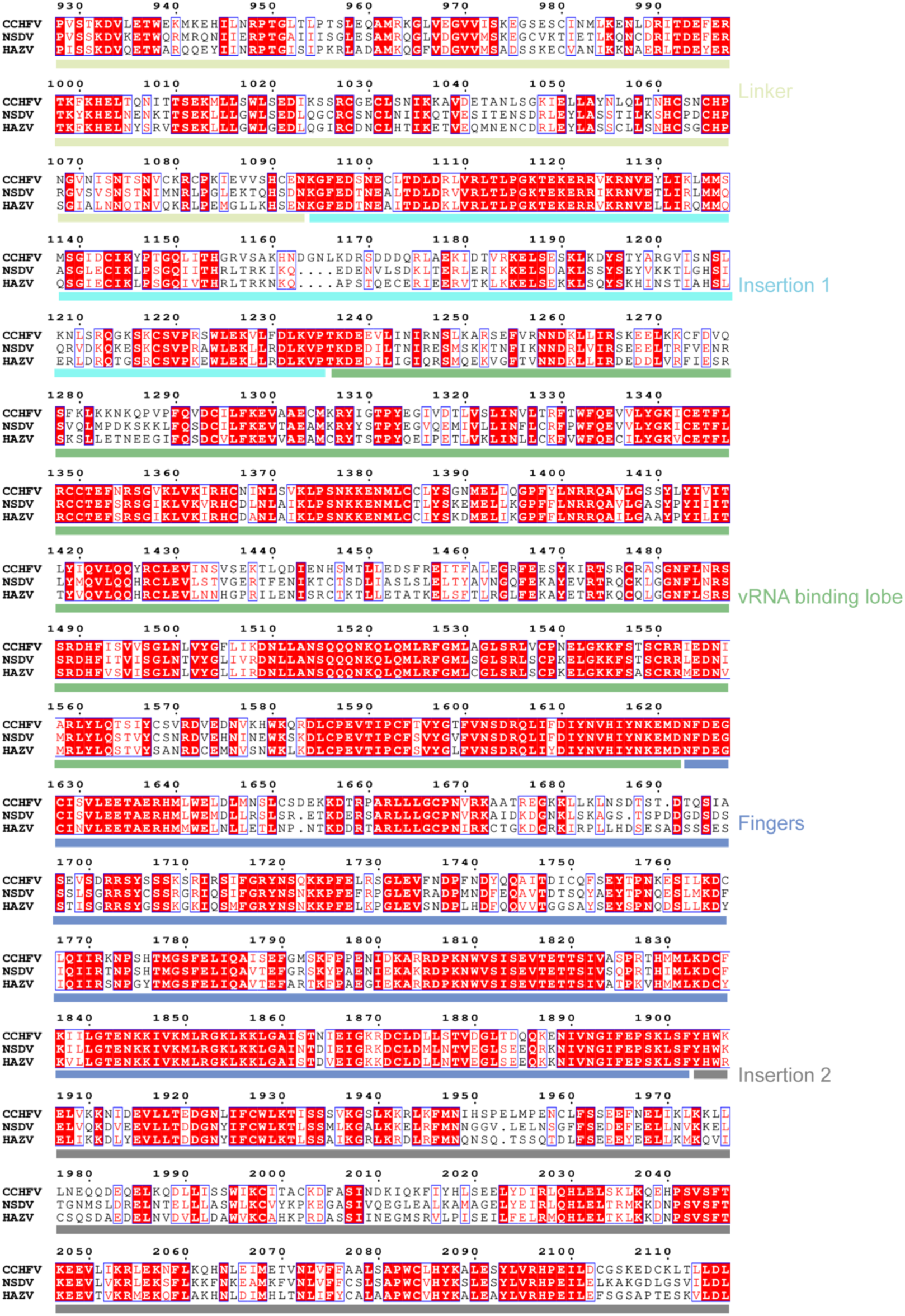
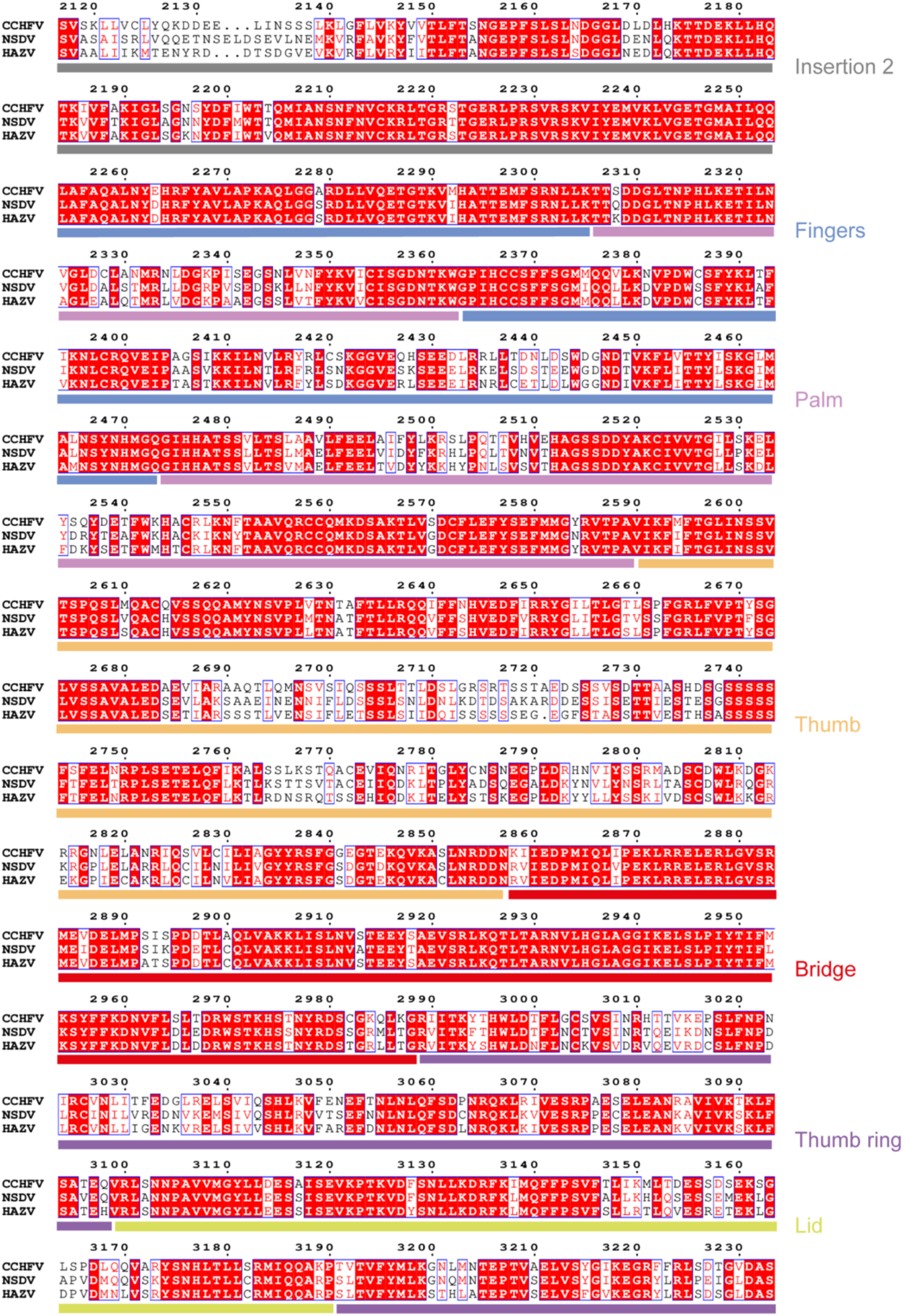
Sequence alignment for nairovirus L-protein core. Sequence alignment for Crimean-Congo Haemorrhagic Fever Virus (Uniprot Q6TQR6), Nairobi sheep disease virus (Uniprot D0PRM7), and Hazara virus (Uniprot A6XA53) L-protein’s. Domains have been annotated above the sequences.

**Supplementary figure 4.**
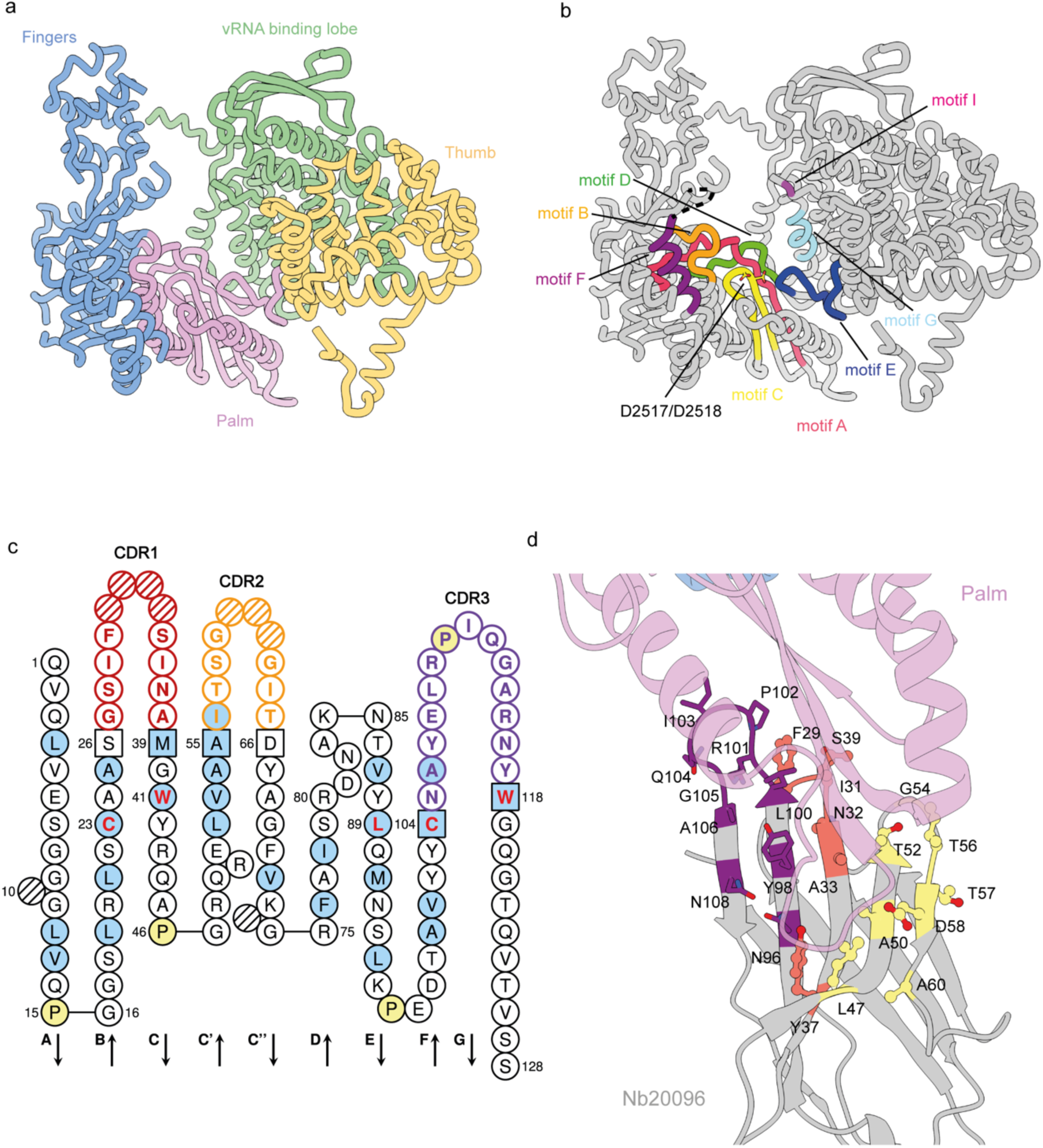
CCHFV-L RNA and NTP binding motifs and Nb20096 binding site. a) The RdRp has been annotated to show the palm (pink), thumb (yellow), vRNA binding lobe (green), and fingers (blue) sub domains. b) Motifs A-G and the RdRp active site residues are highlighted. c) Amino acid sequence and topology of the Nb20096. Complementarity determining regions (CDR) are annotated. d) The interaction site between the CCHFV-L palm and Nb20096 are shown. Residues from Nb20096 that interact with the palm (pink) are annotated and coloured accordingly: CDR1 (orange), CDR2 (yellow), and CDR3 (purple).

**Supplementary figure 5.**
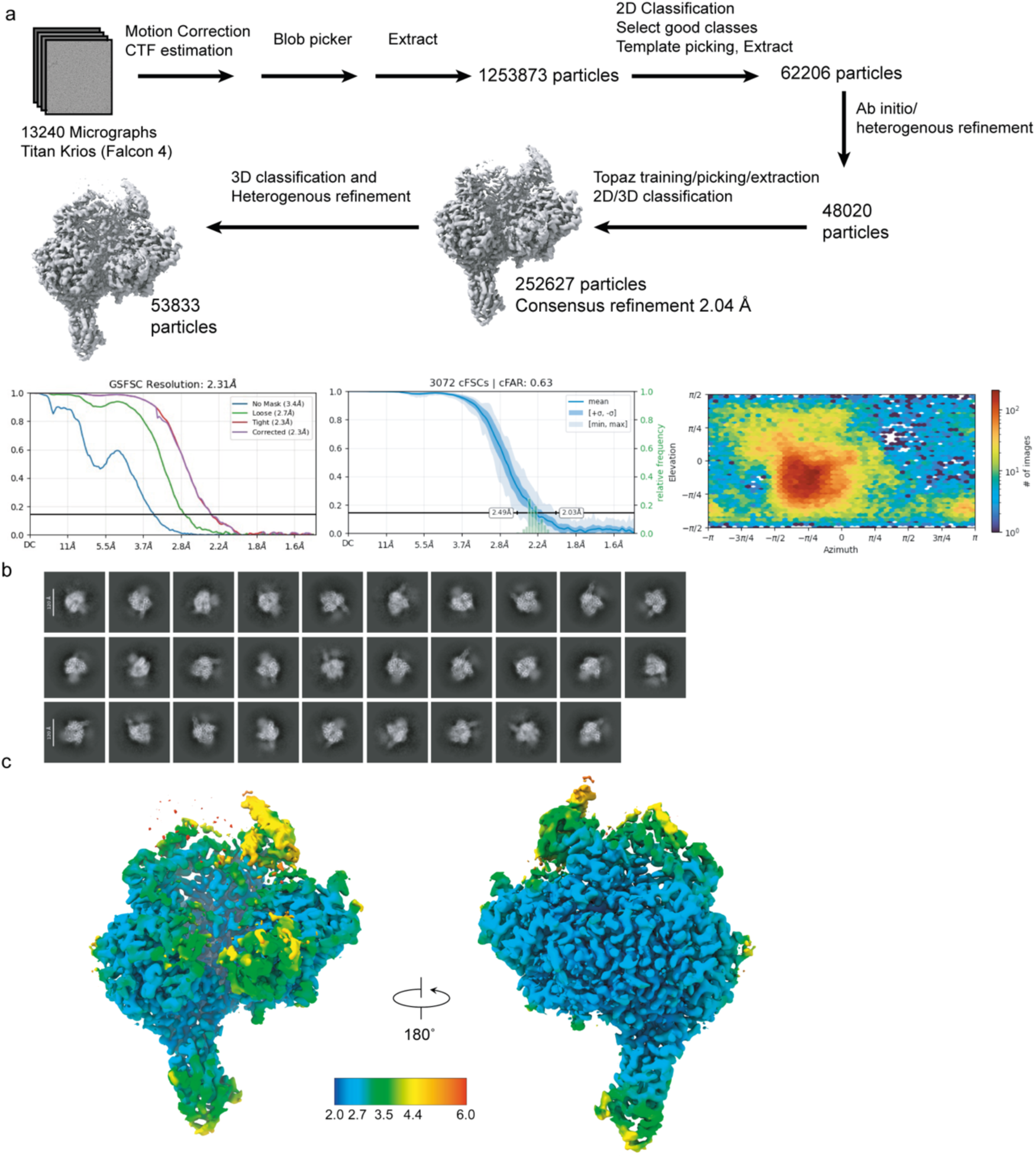
RNA Bound CCHFV-L processing scheme. a) Processing scheme for the RNA free CCHFV-L. b) 2D classes of CCHFV-L. c) Resolution range of the resulting map.

**Supplementary figure 6.**
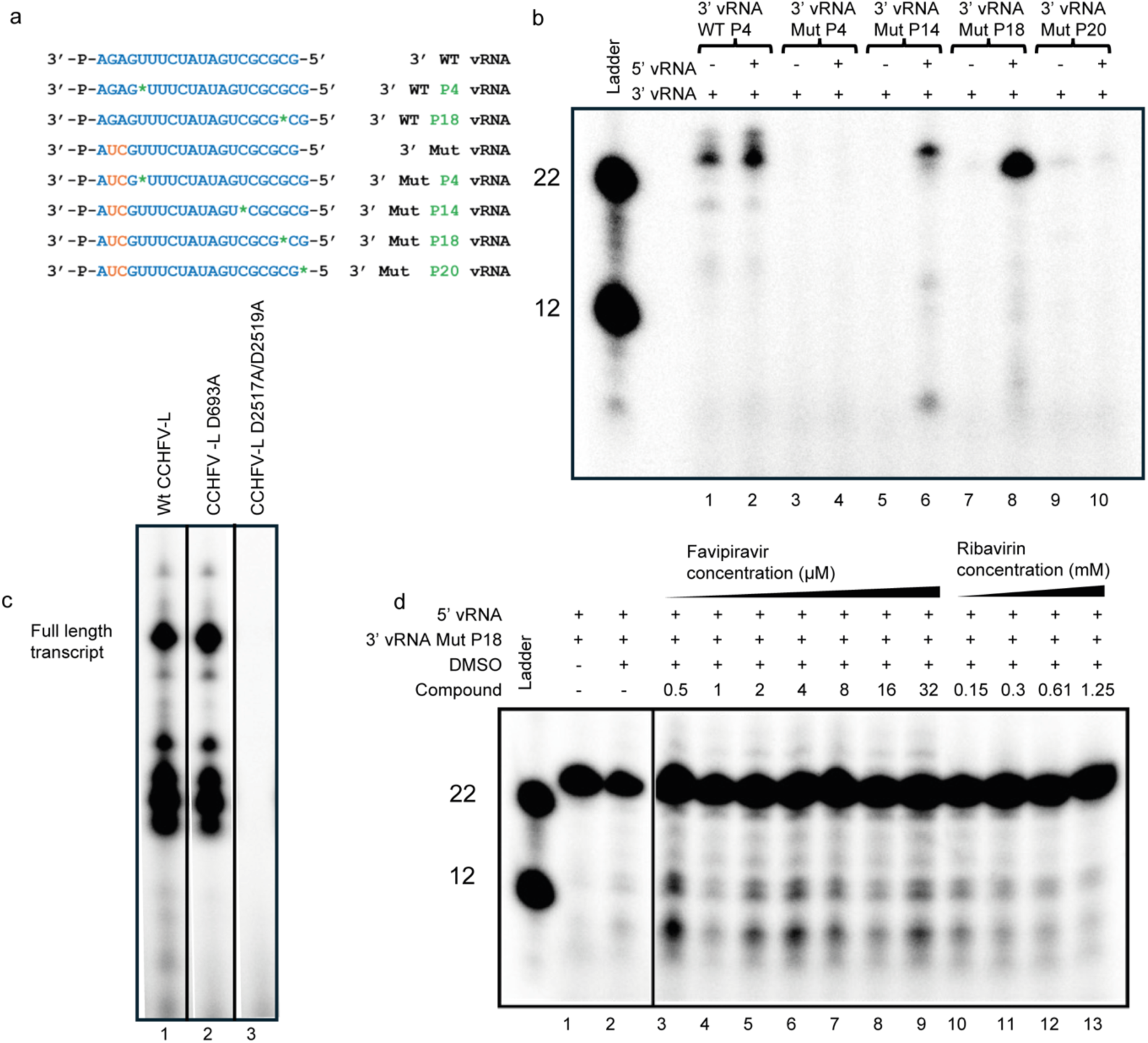
32p Incorporation showing activity on different templates and in the presence of compounds. a) Scheme for 3’ vRNA templates assayed under ^32^P-radiolabelled and fluorescent in vitro conditions. b) Reaction products generated from the fluorescent 3’ vRNAs in the presence and absence of 5’ vRNA. c) Reaction products generated from assays containing 3’ vRNA Mut P18, 5’ vRNA, and 5 nt P1-Cy5 primer using wild-type CCHFV-L, CCHFV-L D693A, and CCHFV-L D2517A/D2518A, respectively. d) Reaction products of ^32^P-radiolabelled assays featuring the 3’ vRNA Mut P18 and 5’ vRNA in the presence of Favipiravir-TP and Ribavirin-TP. Favipiravir-TP concentration titrated from 0.5 to 32 μM, whilst Ribavirin-TP was titrated from 150 μM to 1.25 mM into the extension assays. Compound titration was performed in duplicate with similar results.

**Supplementary figure 7.**
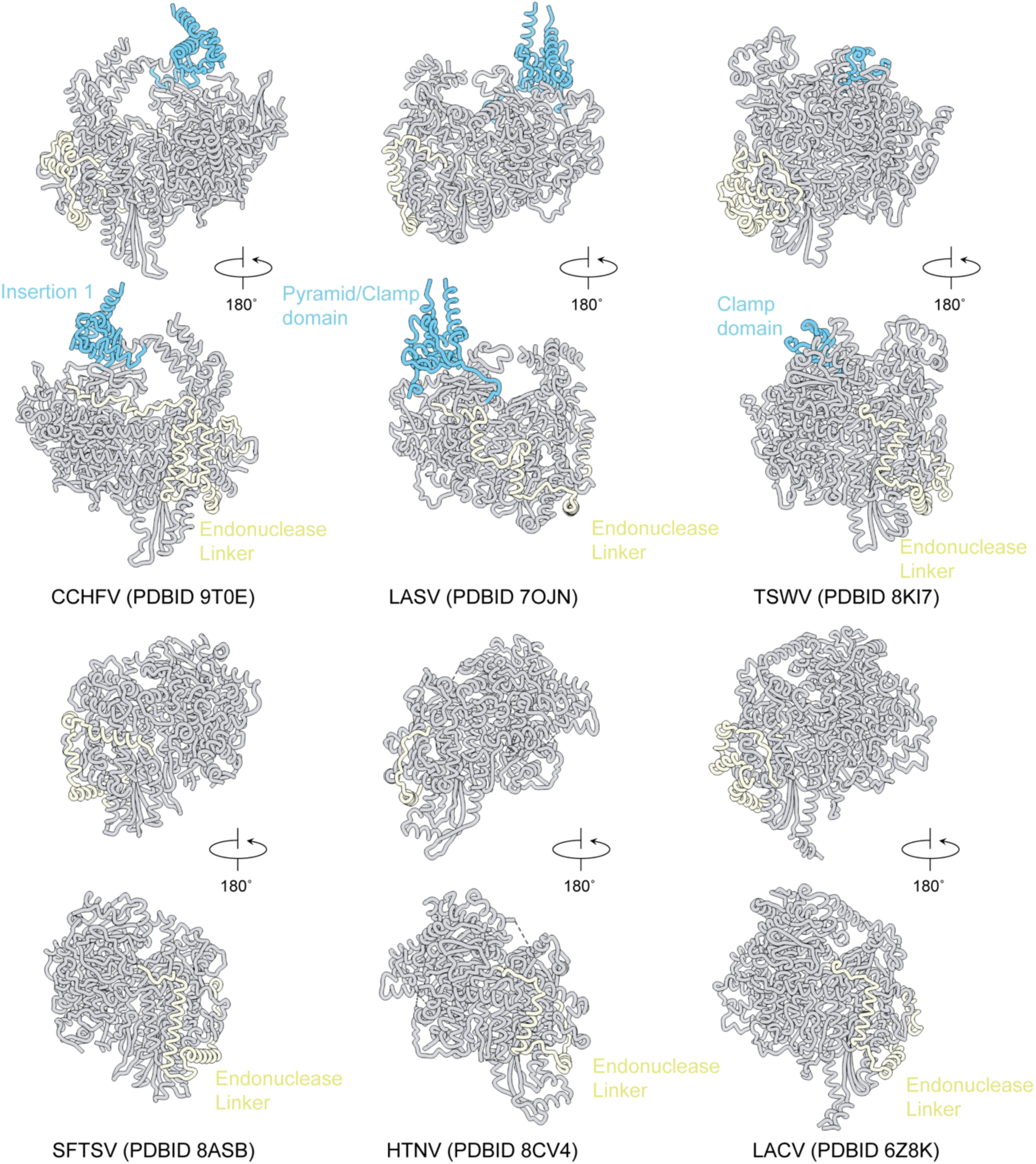
Structural comparison of bunyavirus RdRp. Representatives of the bunyavirus structures which have been determined showing the position of the core RdRp region (grey), insertions/clamp/pyramid domains (cyan), and endonuclease linker (beige). PDB ID’s are annotated in the figure.

